# Structural Studies of Overlapping Dinucleosomes in Solution

**DOI:** 10.1101/753327

**Authors:** A. Matsumoto, M. Sugiyama, Z. Li, A. Martel, L. Porcar, R. Inoue, D. Kato, A. Osakabe, H. Kurumizaka, H. Kono

## Abstract

An overlapping dinucleosome (OLDN) is a structure composed of one hexasome and one octasome and appears to be formed through nucleosome collision promoted by nucleosome remodeling factor(s). In the present study, the solution structure of the OLDN was investigated through integration of small-angle X-ray and neutron scattering (SAXS and SANS, respectively), computer modeling, and molecular dynamics simulations. Starting from the crystal structure, we generated a conformational ensemble based on normal mode analysis, and searched for the conformations that well reproduced the SAXS and SANS scattering curves. We found that inclusion of histone tails, which are not observed in the crystal structure, greatly improved model quality. The obtained structural models suggest that OLDNs adopt a variety of conformations stabilized by histone tails situated at the interface between the hexasome and octasome, simultaneously binding to both the hexasomal and octasomal DNA. In addition, our models define a possible direction for the conformational changes or dynamics, which may provide important information that furthers our understanding of the role of chromatin dynamics in gene regulation.

**Statement of Significance:** Overlapping dinucleosomes (OLDNs) are intermediate structures formed through nucleosome collision promoted by nucleosome remodeling factor(s). To study the solution structure of OLDNs, a structural library containing a wide variety of conformations was prepared though simulations, and the structures that well reproduced the small angle X-ray and neutron scattering data were selected from the library. Simultaneous evaluation of the conformational variation in the global OLDN structures and in the histone tails is difficult using conventional MD simulations. We overcame this problem by combining multiple simulation techniques, and showed the importance of the histone tails for stabilizing the structures of OLDNs in solution.

## Introduction

Nucleosomes are fundamental structural units of chromatin, which enable eukaryotic genomic DNA to be packaged into a nucleus. The canonical nucleosome consists of a histone octamer and about 150 base-pairs of DNA. The histone octamer is composed of two copies each of histones H2A, H2B, H3, and H4, and the DNA segment tightly wraps around its surface (1). For transcription, therefore, the DNA wrapped around the octamer must be unwrapped. This is accomplished by RNA polymerase II, which unwraps the nucleosomal DNA in stepwise fashion during the transcription elongation process (2, 3).

Nucleosomes are dynamic entities that change their position along genomic DNA [e.g. Segal and Widom (4)]. In particular, rearrangement of nucleosome positioning around transcription initiation sites is thought to play a regulatory role in transcription initiation (5). This nucleosome remodeling process is likely mediated by nucleosome remodeling factors (6, 7). It has been reported, for example, that if two nucleosomes are closely positioned, one of the nucleosomes will invade the DNA of its neighbor, probably through nucleosome remodeling, and adopt an unusual structure called an overlapping dinucleosome (OLDN) (8, 9). We previously reconstituted an OLDN and determined its crystal structure (10). Within the OLDN structure, a histone hexasome lacking an H2A-H2B dimer associates with a canonical octasomal nucleosome, and a 250-base-pair DNA segment wraps around the two histone sub-nucleosomal moieties. Sequence mapping using micrococcal nuclease (MNase)-digested HeLa cell chromatin showed that MNase-protected, 250-base-pair DNA segments accumulate in regions just downstream of transcription start sites (10), which suggests OLDNs are formed during the transcription initiation process.

The OLDN structure is asymmetric. To understand the structure and dynamics of OLDNs in solution, we measured its small-angle X-ray and neutron scattering (SAXS and SANS, respectively). The obtained scattering curves were in near agreement with one calculated from the crystal structure, indicating that the asymmetric structure is maintained in solution. However, the bump peak positions were shifted slightly to a lower *Q*, and the observed gyration radius was slightly enlarged. This may reflect structural fluctuation caused by adoption of several stable conformations and/or the lack of histone tails, which could not be observed in the crystal structure.

In the present study, we used computer modeling and molecular dynamics (MD) simulations to model OLDN conformations, including the histone tails, and screened for structures that well-reproduced the experimental data. To do this, we first generated a large number of conformations from the crystal structure by deforming the DNA along the lowest frequency normal modes. We then looked for the conformations that well reproduced the SAXS and SANS data. Although, individually, the SAXS or SANS data were not sufficient to uniquely determine the solution conformations of OLDNs, integration of the SAXS and SANS data prevented the model structures from being overfitted to one or the other data set, which enabled us to successfully narrow the size of the conformational ensemble in solution. Finally, we conducted MD simulations by using each conformation of the ensemble as an initial structure to evaluate the structural stability of OLDNs and investigate their dynamic features in more detail. The results indicate that OLDNs adopt a wide variety of conformations in solution, each of which is stabilized by histone tails situated at the interface of the hexameric and octameric histones. Furthermore, analysis of the conformations can tell us the likely direction of the conformational changes. Such dynamics information may increase our understanding of the assembly and disassembly of OLDNs, which may provide the structural foundation for nucleosome rearrangement within chromatin.

## Materials and methods

### Sample preparation

Recombinant human histone proteins (H2A, H2B, H3.1 and H4) were purified as described previously (11, 12). Histone octamer was reconstituted and then purified using Superdex200 (GE Healthcare) gel filtration column chromatography (11). OLDNs were reconstituted with 250-bp DNA fragments and purified as described previously (10). For SAXS, purified samples were dialyzed against 20 mM Tris-HCl buffer (pH 7.5) containing 50 mM NaCl, 0.1 mM MgCl_2_, and 1 mM dithiothreitol. For SANS, purified samples were dialyzed against 20 mM Tris-HCl (pH 7.5) buffer containing 50 mM NaCl, 0.2 mM MgCl_2_, and 1 mM dithiothreitol with different amounts of D_2_O (0, 40, 65 and 100%, respectively). The concentration of OLDNs was calculated from the absorbance by the DNA (260 nm) and determined as the DNA-histone 14-mer complex (1.00 mg/ml DNA corresponds to 2.26 mg/ml complex).

### Solution Scattering

SAXS and SANS were conducted to observe the structure of OLDN in aqueous solution: SAXS was used to examine the overall shape of OLDNs, while SANS was employed to separately observe the structures of the elements comprising OLDNs, including the histone domains and the DNA.

SAXS experiments were performed with a SAXS camera installed at BL10C of the KEK Photon Factory (Ibaraki, Japan). Using a two-dimensional semiconductor detector (PILATUS3 2M), SAXS intensity was measured for 300 s using a time slice of 15 s by checking the radiation damage on each sample. The covered *Q*-range was from 0.008 Å^−1^ to 0.25 Å^−1^: *Q*=(4_*π*_/*λ*)sin(*θ*/2), where *λ* and *θ* are the wavelength of the incident beam and the scattering angle, respectively. After checking the radiation damage, the SAXS pattern was converted to a one-dimensional scattering profile, after which standard corrections for the initial beam intensity, background scattering, and buffer scattering were applied. Finally, the obtained SAXS intensity of the sample was normalized to the absolute scale using a glassy carbon standard. The samples were solutions of the DNA-histone 14-mer complexes in buffer at a concentration of 0.5 or 3.0 mg/ml. No particle interference was observed with either solution.

SANS experiments were performed on D22 installed at the High Flux Reactor of the Institut Laue-Langevin. To cover the *Q* -range from 0.008Å^−1^ to 0.25 Å^−1^, the SANS intensity was measured at two sample-to-detector distances (SDDs), 5.6 m and 2.0 m, using a 6-Å neutron beam. The measured two-dimensional scattering pattern was converted to a one-dimensional scattering profile of solute by following the standard procedure, circular averaging, correction of transmission, and substations of buffer scattering and background. Thereafter, the SANS profiles with SDDs of 5.6 m and 2.0 m were merged into one profile using GRASP software (http://www.ill.eu/instruments-support/instruments-groups/groups/lss/grasp/home/).

For small-angle scattering, scattering intensity *I*(*Q*) is described as follows,

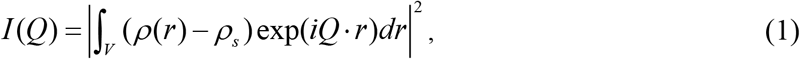

where *V*, ρ and ρ_s_ are the volume of the solute and the scattering length densities of the solute and solvent, respectively. With neutron scattering, there is an isotope effect on scattering length, which is especially large between hydrogen (−3.74 fm) and deuterium (+6.67 fm). Reflecting this difference, ρ_s_ can be tuned by mixing H_2_O and D_2_O to a proper ratio. This is called the “contrast variation technique.” As shown in Fig. S1 in the Supporting Material, the scattering length densities of histone and DNA are matched to those of 40% D_2_O and 65% D_2_O, respectively. Following Eqn.1, this means that in 40% D_2_O solution a histone is invisible, and only the structures of the DNA in OLDNs can be observed; in a same manner, the structures of only histones in OLDN can be observed in 65% D_2_O solution. Using this approach, we measured the SANS profiles of OLDNs (3 mg/ml DNA-histone 14-mer complex) in solutions containing 0%, 40%, 65% and 100% D_2_O.

### DNA treated at base-pair step level

One of the authors developed a method for studying the static and dynamic structures of double-stranded DNA using the base-pair step parameters (Tilt, Roll, Twist, Shift, Slide, Rise) as internal coordinates (13–15). In the present study, we used this method to model the missing base-pairs of the DNA in the X-ray crystal structure and to deform the DNA.

With this method, base-paired residues in double-stranded DNA are treated as a rigid body, and the relative position and orientation of two adjacent rigid bodies (or base-pairs) are described in terms of six base-pair step parameters (Tilt, Roll, Twist, Shift, Slide, Rise). Deviation of the geometry from the equilibrium increases the conformational energy (dimer step energy) *E*_*d*_ described as

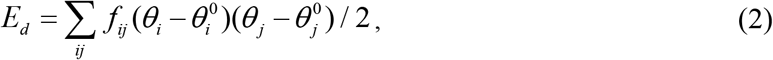

where *θ*_*i*_ and 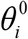 are the instantaneous and equilibrium values of the base-pair step parameters, and *f*_*ij*_ is the force constant. Olson et al.(16) derived these values (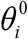 and *f*_*ij*_) in a sequence-dependent manner – i.e., different constants for different kinds of dimer steps – by analyzing a large number of crystal structures. In the present study, we used those constants unless otherwise noted. The total conformational energy of the double-stranded DNA is described as ∑*E*_*d*_.

### Deformation of OLDN by changing the DNA conformation

We deformed the crystal structure of OLDN by changing the conformation of the DNA while the structures of the histone octamer and hexamer were fixed. Because the base-pair step parameters were used as the internal coordinates, each conformation of the DNA was described by a 6×(*N*−1)-dimensional vector Θ, where *N* is the number of base pairs. The deformation of the DNA was performed by changing the base-pair step parameters along the normal mode vectors as

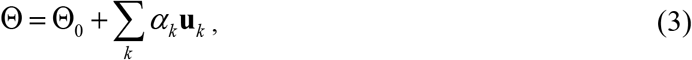

where Θ_0_ describes the crystal structure, *α*_*k*_ is the magnitude of the deformation, and **u**_*k*_ is the *k*-th lowest-frequency normal mode vector. In the present study, we used the five lowest frequency modes (*k*=1,2,3,4,5) in the summation. It should be noted that different conformations are obtained by giving different sets of numbers (*α*_1_,*α*_2_, *α*_3_,*α*_4_, *α*_5_).

The normal mode vectors were obtained using a computational procedure that was nearly the same as that used for linear DNA (15). The differences were that, in the present computation, we assumed that the crystal structure was in the minimum energy conformation – i.e., the values of the base-pair step parameters in the crystal structure were used as the equilibrium values 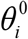 in Eqn. 2 – and that we used very large force constants *f*_*ij*_ (100 times larger than those derived by Olson et al. (16)) for the DNA wrapped around the histone core proteins so that they would not change their conformations easily. We assigned normal force constants only to the base-pair steps (chain I:129-152, chain J:99-122) in the linker DNA region, where the DNA did not wrap around the core proteins. To build the deformed atomic model of an OLDN, after deformation of the DNA, the histone octamer and hexamer were put into the same geometry as in the crystal structure with respect to the wrapping DNA.

### Modeling OLDNs with histone tails

The crystal structure of the OLDN lacked the histone tails. However, to reproduce the experimental SAXS profile well, we found it necessary to include the histone tails in the models. We therefore modeled OLDNs with histone tails in the following way. We started from the structure with the minimum χ^2^ for the SAXS profile, which was obtained by deforming the crystal structure without the histone tails. Based on this structure, the initial histone tail conformations were modeled using the program MODELLER (17, 18). We then performed 107 independent MD simulations using the simulated annealing method for OLDN with the modeled histone tails to obtain distinct tail structures. The model was put in a box with dimension of 26.4×2.4×4.6 nm^3^, which was kept unchanged during the simulations (constant volume). The box was filled with the TIP3P water containing Na^+^ and Cl^−^ ions to neutralize the system and maintain the salt concentration at 150 mM. The simulations started with a rapid increase of the temperature from 3 K to 1500 K in 0.15 ns, followed by a more gradual reduction of the temperature to 0 K in 1.5 ns. In all the simulations, only the modeled histone tails were allowed to move, while the DNA and histone core proteins were restrained. Different initial velocities to the atoms were assigned in different simulation runs to obtained different tail structures. The simulation was carried out using GROMACS (19–25) with Amber 14sb+bsc1 force field (26) to describe the nucleosome. The temperature was controlled using the V-rescale method (27). The final conformation in each simulation run was collected. From the 107 collected models with different conformations of the histone tails, we selected 50 whose histone tails were bound to the nucleosomes and were not extended outward.

Ideally, the above computations should have been performed with all the different OLDN conformations built by deforming the crystal structure. That was not possible, however, due to the computational time and resources it would have required. Instead, we replaced the histone core proteins lacking histone tails with proteins with tails in the aforementioned 50 models. This enabled us to build 50 different conformations of OLDNs with histone tails from the model without them.

### Selection scheme

To select appropriate atomic models that well reproduced the experimental profiles, we used the χ^2^ value defined as

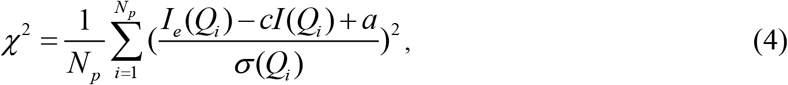

where *N*_*p*_ was the number of experimental points *Q*_*i*_; *I*_*e*_(*Q*_*i*_), and *I*(*Q*_*i*_) were the experimental and computed profiles, respectively; *σ*(*Q*_*i*_) was the experiment error; *c* was a scale) factor given by

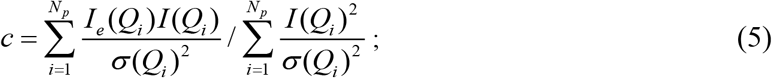

and *a* was the offset that accounts for possible systematic errors due to mismatched buffers in the experimental data (28). Smaller χ^2^ values indicated a better fit to the experimental profile.

### Conformational analysis using six rigid-body parameters

To describe the global conformation of each OLDN model, six rigid-body parameters (*dX*, *dY*, *dZ*, *dρ*_*X*_, *dρ*_*Y*_, *dρ*_*Z*_), which reflected the positions and orientations of the two nucleosomes relative to each other, were computed as follows. First, we defined the reference coordinate system on the X-ray crystal structure of a mononucleosome (PDB: 3LZ0 (29)). The origin was set on the center of mass, and the xyz-axes were defined by the principal axes of inertia (Fig. 1a). Note that we included both the DNA and proteins in the calculation. The z-axis appeared to coincide with the superhelical axis of the DNA, and the y-axis appeared close to the dyad symmetry axis. Then, the histone core proteins of the mononucleosome were fitted to the corresponding proteins (RMS-fitting) in the OLDN model. Through this fitting, the origin and xyz-axes were defined locally in each nucleosome (Fig.1b). The xyz-axes were described by a rotation matrix, **R**_*i*_, and the origin was described by a vector, **o**_*i*_. The subscript *i* was used to differentiate the two nucleosomes within an OLDN. We assumed that **R**_1_ and **o**_1_ were for the octasome, and **R**_2_ and **o**_2_ were for the hexasome. Finally, using the CEHS scheme (30, 31) to compute the base-pair step parameters, the six rigid-body parameters were computed. The angular parameters (*dρ*_*X*_, *dρ*_*Y*_, *dρ*_*Z*_) were computed from the rotation matrices **R**_1_ and **R**_2_. The translational parameters (*dX*, *dY*, *dZ*) were computed as 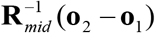, where **R**_*mid*_ described the “middle frame” between **R**_1_ and **R**_2_. The parameters *dρ*_*X*_, *dρ*_*Y*_, and *dρ*_*Z*_ respectively correspond to the base-pair step parameters Tilt, Roll, and Twist, while *dX*, *dY*, and *dZ* correspond to Shift, Slide, and Rise.

**Figure 1.**
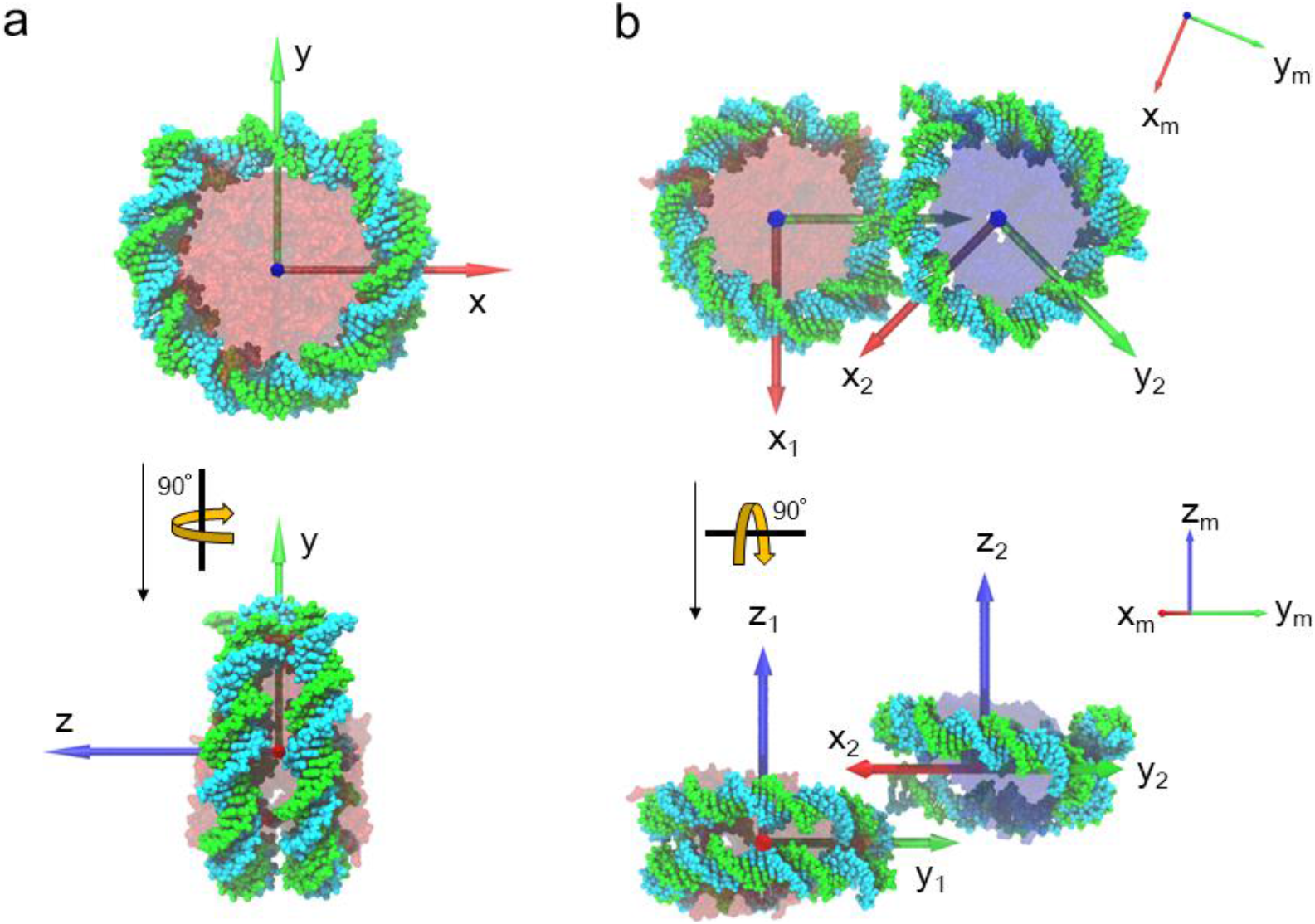
(a) The xyz-coordinate system defined on the reference atomic model. The x-(red arrow), y-(green), and z-axes (blue) were defined by the principal axes of inertia of the X-ray crystal structure of the mononucleosome (PDB: 3LZ0). The origin was set on the center of mass of the structure. (b) For both the octasome and hexasome, the xyz-coordinate system was defined (x_1_y_1_z_1_ and x_2_y_2_z_2_, respectively) by fitting the X-ray crystal structure of the mononucleosome shown in (a). For easy understanding of the six rigid-body parameters, the octasome and hexasome are situated such that the z_1_− and z_2_− axes are parallel, and the angle between the x_1_− and x_2_− axes is 45 degree, ignoring the connection between the two nucleosomes. Transparent red: octameric proteins, transparent blue: hexameric proteins. The middle frame (x_m_y_m_z_m_) is also shown.

### MD simulations for model verification

We performed MD simulations with the models that well reproduced the experimental SAXS and SANS profiles to see whether they existed stably in solution. As described above, the histone tails in these models were built for the specific structure with the smallest χ^2^ for the SAXS profile. Here, we optimized the conformations of the histone tails for each model before each simulation run as follows. First, the histone tails were replaced with the extended tails modeled using the program MODELLER. We then performed MD simulations using the simulated annealing technique to enhance the conformational changes in the histone tails, while the rest of the structure remained fixed. The same procedure was applied when we added the histone tails to the model without tails. In that case, we repeated the procedure more than 100 times with the same model to build a variety of conformations of the histone tails. This time, we performed the simulation only once for each model. After remodeling the histone tails, we performed conventional MD simulations (NVT) at a temperature of 300 K with no restraint using GROMACS with the Amber 14sb+bsc1 force field. The temperature was controlled using the V-rescale method.

### Results and Discussion

#### Solution scattering

Figures 2a and 2b show SAXS profiles in an aqueous solution and their Guinier plots (log *I*(*Q*) vs *Q*^2^). Through least square fitting at low *Q* with the Guinier formula 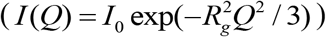, where *R*_*g*_ is the radius of gyration (Fig. 2b), the radius of gyration was calculated to be 57.5±0.3 Å. In Fig. 2a, the SAXS profile calculated from the crystallographic data (PDB: 5GSE) is indicated by a blue line. The experimental and computational SAXS profiles were similar, but the peak positions in the experimental profile were shifted slightly to a lower *Q*. This indicates that, as a whole, the structure in aqueous solution is basically the same as that in crystal, but there is a slight structural modulation. The radius of gyration of the crystal structure was calculated to be 46.3 Å (Table 1), which is smaller than that obtained with SAXS or SANS. It should be noted that calculation of *R*_*g*_ for the crystal structure included no contribution from the missing histone tails, which were not observable.

**Table 1.** Radii of gyration.

|  | Solution<br>D<sub>2</sub>O conc. | Experiment | Crystal |
| --- | --- | --- | --- |
| SAXS | 0 % | $57.5 \pm 0.3$ Å | 46.3 Å |
| SANS | 0 % | $55.3 \pm 0.6$ Å | 46.4 Å |
| | 40 % | $61.1 \pm 1.6$ Å | 51.6 Å |
| | 65 % | $50.1 \pm 0.4$ Å | 39.3 Å |
| | 100 % | $54.8 \pm 0.4$ Å | 43.4 Å |

**Table 1.** Radii of gyration.
|  | Solution | Experiment | Crystal |
| --- | --- | --- | --- |
|  | D<sub>2</sub>O conc. |  |  |
| SAXS | 0 % | 57.5 ±0.3 Å | 46.3 Å |
| SANS | 0 % | 55.3 ±0.6 Å | 46.4 Å |
| | 40 % | 61.1 ±1.6 Å | 51.6 Å |
| | 65 % | 50.1 ±0.4 Å | 39.3 Å |
| | 100 % | 54.8 ±0.4 Å | 43.4 Å |

**Figure 2.**
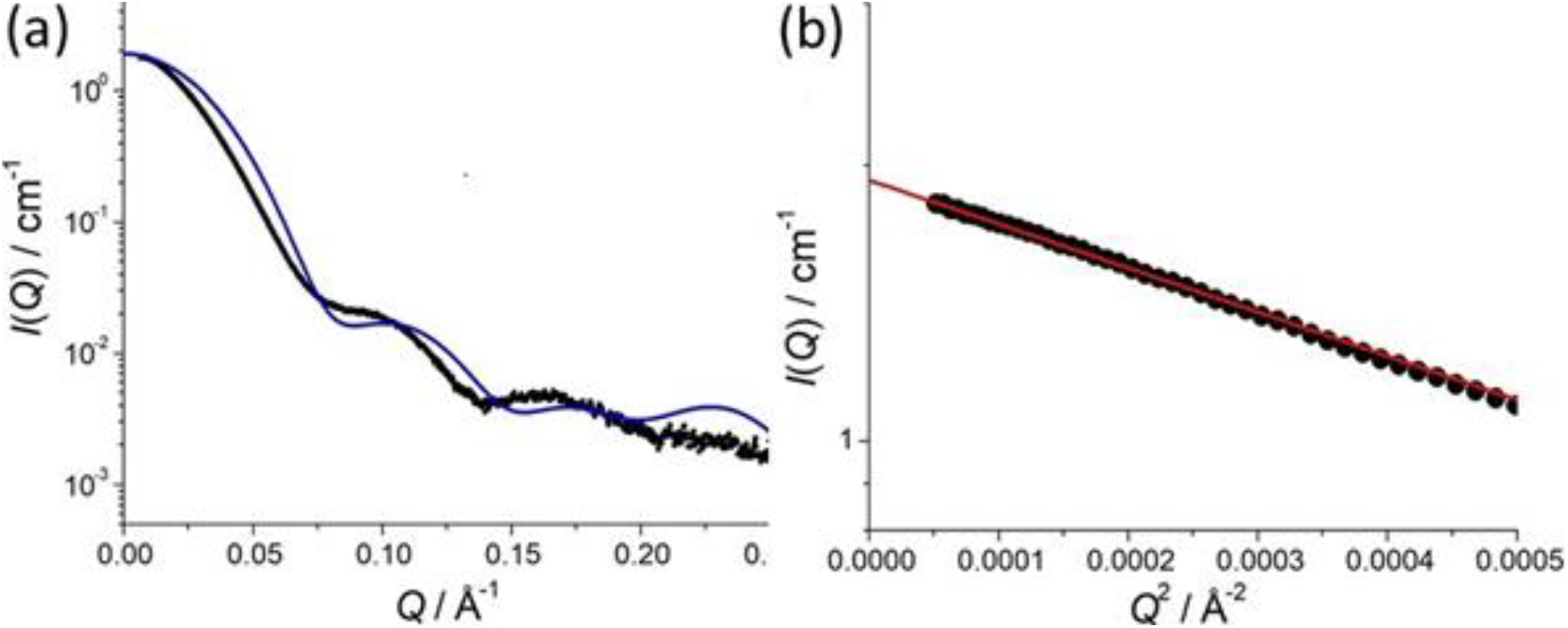
(a) SAXS profiles for OLDNs. Black dots show the experimental SAXS profile, and a blue line indicates the calculated profile based on the crystal structure (PDB: 5GSE). (b) Guinier plots. Filled circles are the experimental results, and the red line shows the result of the least square fitting with the Guinier formula (see text).

To examine the structure of OLDN in more detail, we conducted contrast-variation SANS (CV-SANS) measurements, which provide structural information about the histones and DNA within OLDNs separately (see Figure S1). Figure 3 shows the SANS profiles with their Guinier plots as insets for OLDNs in 0%, 40%, 65%, and 100% D_2_O. The radii of gyration are listed in Table 1. The SANS profiles in 40% and 65% D_2_O respectively correspond to the profiles for the DNA and histones within OLDNs. As expected, therefore, the radius of gyration in 40% D_2_O was the largest and that in 65% which indicates that the DNA twines were wrapping around histone cores. Interestingly, even though the radii of gyration were larger than those calculated from the crystal structure in all the contrast conditions, the SANS profiles were becoming more similar in the higher *Q*-region (roughly *Q* > 0.10 Å^−1^). This suggests that although the individual nucleosomes, hexasomes, and octasomes, have basically the same structures in aqueous solution as in crystal, they are able to adopt dynamic configurations. To elucidate these molecular structures at the atomic level, we constructed structural models by integrating the SAXS and CV-SANS experiments with the computational methods. In this modeling, the DNA and histone tails missing from the crystallographic data were explicitly considered.

**Figure 3.**
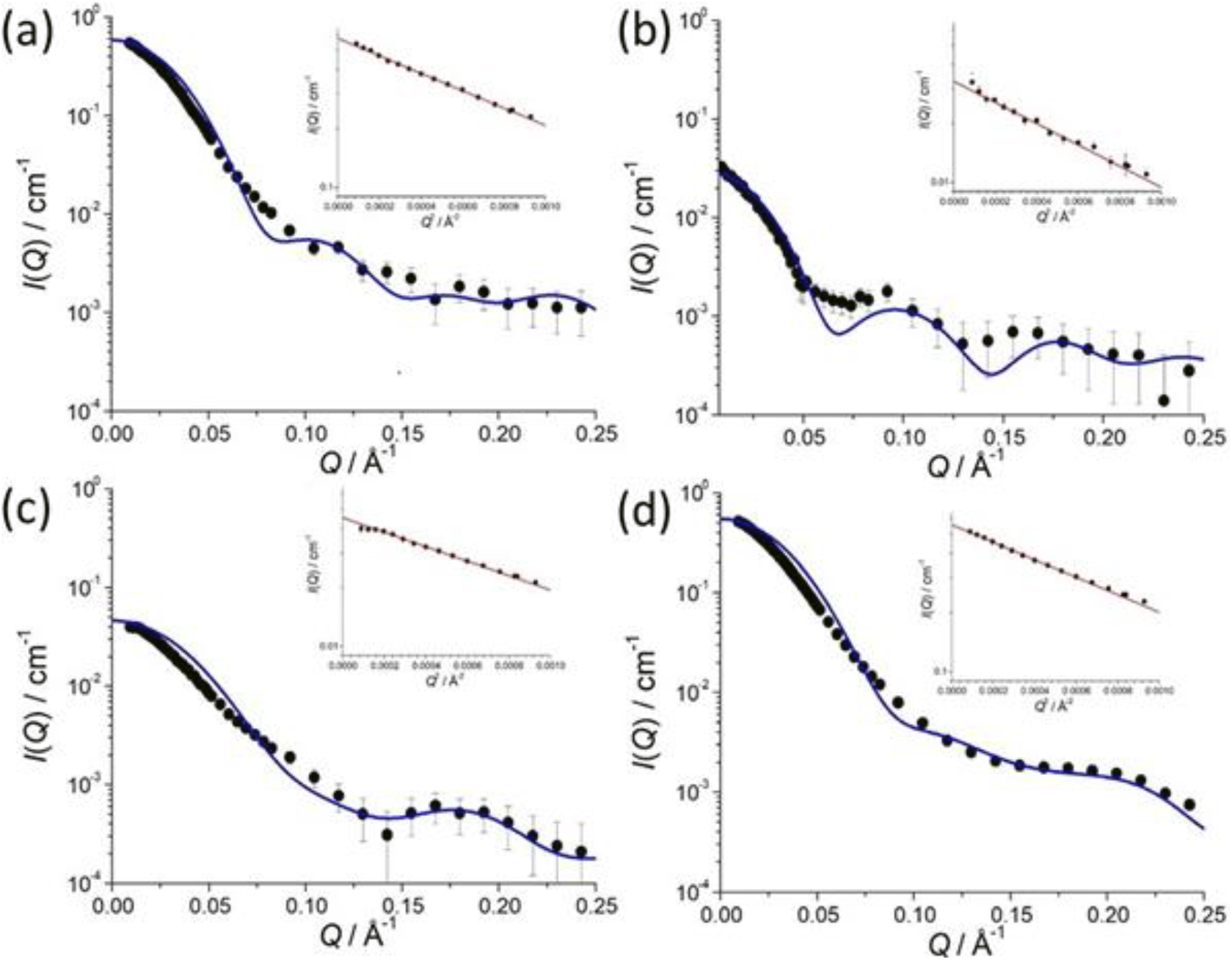
CV-SANS profiles and their Guinier plots. (a) 0% D_2_O (H_2_O), (b) 40% D_2_O, (c) 65% D_2_O, (d) 100% D_2_O. Blue lines show the SANS profiles calculated based on the crystal structure (PDB: 5GSE). Insets are Guinier plots in which red lines show the results of the least square fitting with the Guinier formula (see text).

#### Overview of the construction of atomic models consistent with the scattering data

The computational procedure we used to obtain atomic models consistent with the scattering data and to investigate OLDN dynamics is outlined in Fig. 4. It consisted of six steps and started from the OLDN crystal structure (PDBID 5GSE (10)). Here, we provide an overview of each step. The steps are described in detail in the Supplement and in the Materials and Methods.

**Figure 4.**
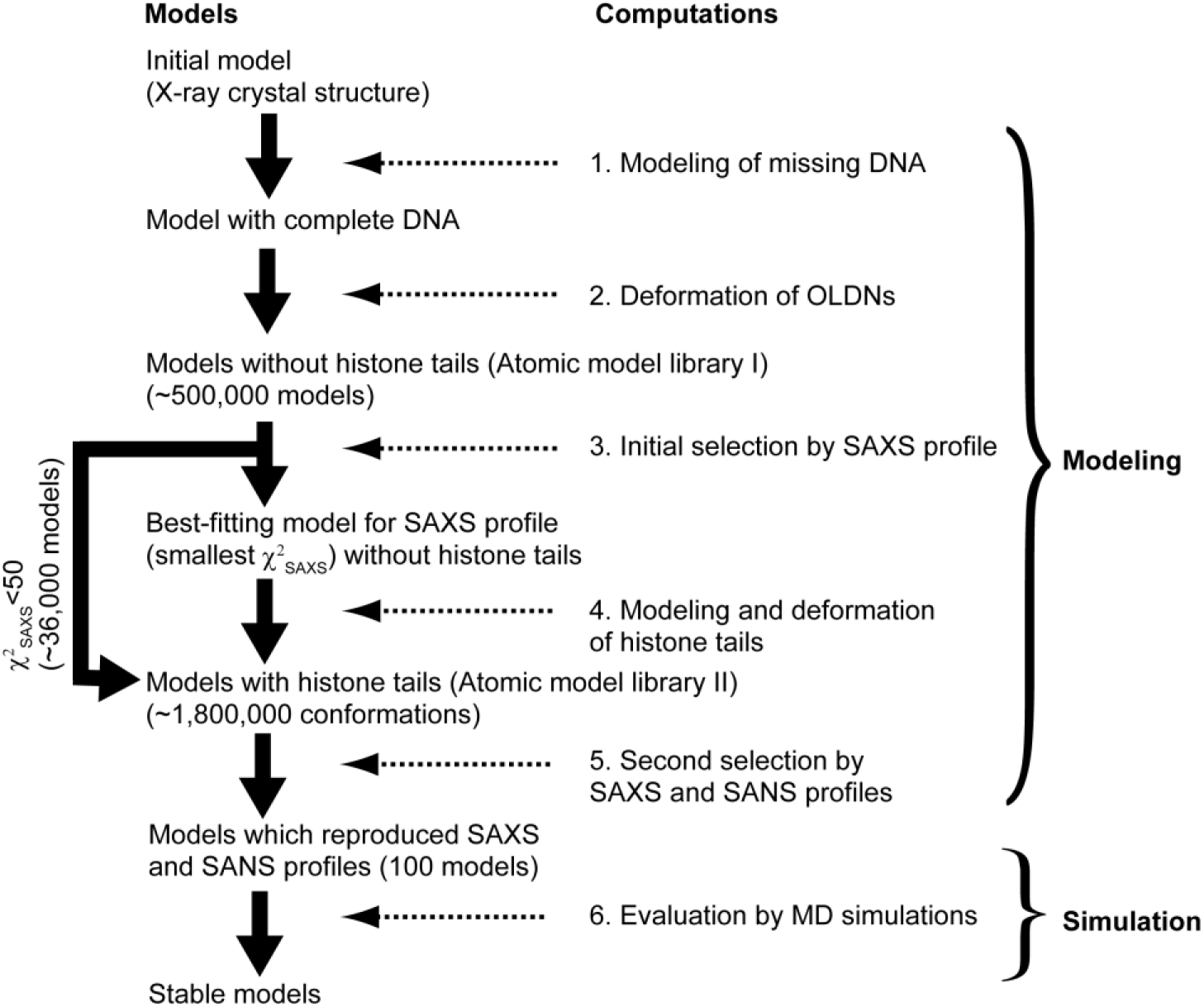
Overview of the computations performed in this study.

##### Step 1: Modeling of the missing DNA

The X-ray crystal structure of an OLDN lacks five successive base pairs (chain I:131-135 and chain J:116-120). To model these base pairs (i.e., to fill the gap), we considered a DNA model with seven base pairs that had the same sequence as the missing base-pairs plus two adjacent base pairs (chain I:130-136, chain J:115-121). We then minimized the total conformational energy of the DNA model, which included a penalty function to force the base pairs on both ends to have the same geometry as in the crystal structure. The resultant DNA model was inserted into the crystal structure by fitting the end base pairs to the corresponding ones in the crystal structure (RMS fitting). Hereafter, we will refer to this X-ray crystal structure with the DNA gap filled by the model as simply the crystal structure.

##### Step 2: Deformation of OLDNs

To construct the atomic model of OLDNs fitted to the SAXS and SANS data, we prepared a large number of different conformations. For this purpose, we first generated a wide variety of different DNA conformations by deforming the crystal structure using the five lowest-frequency normal modes (see Materials and Methods). Then, the histone octamer and hexamer, whose structures were fixed as the crystallographic structures, were put in the same geometry with respect to the wrapping DNA in the crystal structure. It should be noted that we assumed that the crystal structure was the minimum energy conformation and allowed only the linker DNA (chain I:129-152, chain J:99-122) to change its conformation. With this approach, we produced more than 500,000 atomic models of an OLDN (the atomic model library I), which differed from one another by at least 2 Å in RMSD (Root Mean Square Deviation).

##### Step 3: Initial selection by SAXS profile

When we calculated χ^2^ for each of the models in library I against the SAXS profile, the minimum χ^2^ was 11, which we considered too large for the model to well reproduce the experimental profile. This may have been due to the lack of the histone tails in the crystal structure. In fact, the tails occupy about 20% to 25% of the total weight of the histones and so cannot be ignored in terms of scattering intensity (32). Therefore, for further analysis, we selected ∼36,000 OLDN models with a relatively small χ^2^ (<50) in which the histone tails were to be modeled.

##### Step 4: Modeling and deformation of histone tails

The initial conformations of the histone tail were prepared using MODELLER (17, 18) based on the OLDN structure with the minimum χ^2^ obtained at step 3. Because histone tails are highly flexible, using GROMACS (19–25) we repeated independent, simulated annealing MD simulations of this OLDN about 100 times, starting from the same initial conformation. In the simulations, only the modeled histone tails were allowed to move; the DNA and histone core regions were restrained. From the ∼100 final models with different tail conformations, we selected 50 whose histone tails were bound and not extending outward (see Fig. S8). We then replaced the histone proteins without tails in the 36,000 models selected at step 3 with the proteins with tails in one of the 50 models through RMS-fitting of the histone core atoms. As a result, we built a new structure library with 1.8 million (36,000×50) different conformations (atomic model library II).

##### Step 5: Second selection of structure with SAXS and SANS

Because the histone tails are very flexible, it is reasonable to consider their multiple conformations – i.e., we considered that in each OLDN model, the histone tails had 50 different conformations. To examine the reproducibility of the models against the experimental SAXS and SANS profiles, we used the mean value of χ^2^ over the 50 conformations, which differed only in the histone tails. Hereafter, we will denote the mean value of χ^2^ as <χ^2^>.

##### Selection using SAXS profiles

We calculated 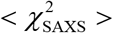(<χ^2^> for the SAXS profile) for all the models in the new library II, where the models have tails. The minimum value of 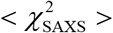 was 5.7, showing that addition of the histone tails improved χ^2^. Indeed, 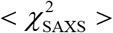 for most of the models was smaller after the histone tails were added (Fig. S7). In Fig. S9, two computed SAXS profiles were compared with the experimental profiles. The solid red line is the profile for the model with the smallest χ^2^ when the histone tails were not considered, and the cyan dashed line is for the model with the smallest 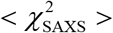. The latter was obtained by averaging 50 different profiles for the model in which multiple conformations of the histone tails were considered. In the lower region of *q* (<0.1 Å^−1^), both profiles well fit to the experimental profile, indicating that the overall shape of OLDN is reproduced, even by the models without the histone tails. However, in the higher region of *q*, deviation of the former from the experimental profile was apparent, demonstrating that addition of the histone tails improved the reproduction of local OLDN structures.

##### Selection using SANS profiles

SANS profiles in 40% and 65% D_2_O give information about the conformations of the DNA and proteins, respectively. We used these two different profiles as a kind of “low-pass filter” to exclude the models with 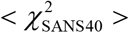 or 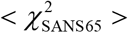 (<χ^2^> for the SANS profile in 40% and 65% D_2_O) higher than given threshold values. By applying these filters to the models with small 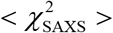, we further limited them by excluding those that satisfied the SAXS profile but not the SANS profiles reflecting the conformations of the DNA or histones. The threshold values were set so that a quarter of the models with 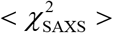 were blocked by each filter (∼1.90 for both 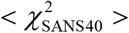 and 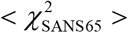). Using these filters, 320 models were extracted from the 500 models with the smallest 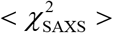 (<∼7.5).

We analyzed the conformations of the models using the six rigid-body parameters, which described the positions and orientations of the two nucleosomes relative to each other within the OLDN (Fig. 1). Figure 5 shows the distributions of the parameters for the 500 models (black open bar) as well as the 320 extracted models (green open bar) (left y-axis). The most significant difference between the two distributions was observed in *dX*. The models with relatively large *dX*(>∼20Å) were excluded by the SANS filters. It should be noted that application of the SANS profile filters in 0% and 100% D_2_O did not noticeably change the distributions (data not shown). This is reasonable because these profiles included the scattering from both proteins and DNA as the SAXS profile, so they should be essentially the same as the SAXS profile.

**Figure 5.**
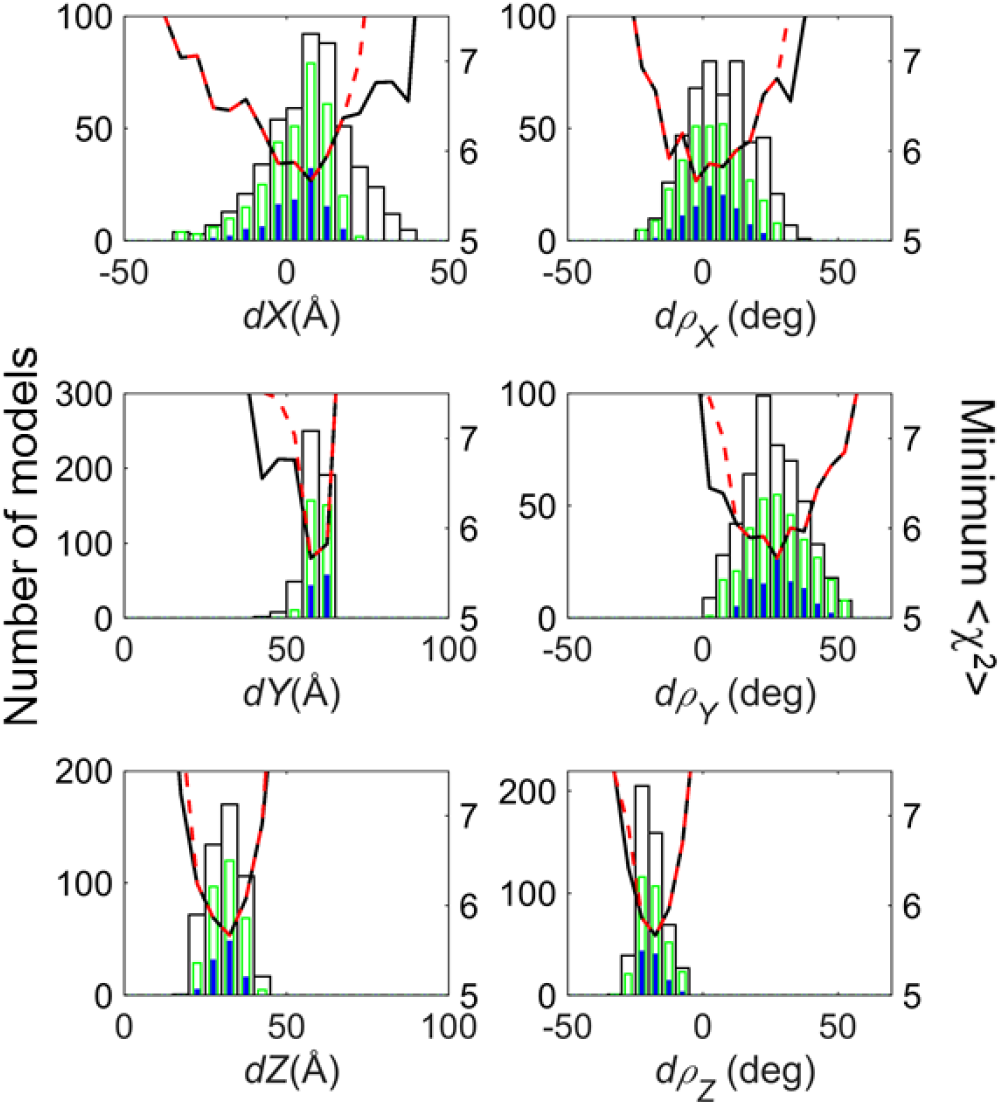
Distributions of the six rigid-body parameters for the atomic models of OLDNs in which multiple conformations of the histone tails are considered (left y-axis). For the 500 models with the smallest 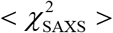, the distributions are denoted by open ck bars. Among the 500 models, the 320 models in which both 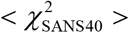 and 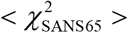 smaller than the threshold values are denoted by open green bars. Among the 320 models, the 100 atomic models with the smallest 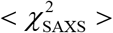 were used as the initial conformations in MD simulations and are denoted by filled blue bars. The bin size was set to 5 (Å or degrees) in all the distributions. The smallest 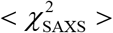 of the models in each bin is also plotted (right y-axis) as a black solid line for the models included in the open black bar and as a red dashed line for models included in the open green bar.

The smallest 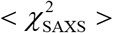 of the models included in each bin is also plotted in Fig. 5. The shape of the plot was similar to that of the distribution of models if inverted, and the conformation with the lowest 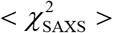 was nearly always located in the highest bin of the distribution. This suggests that a group of models with relatively small 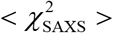 were distributed around the conformation with the smallest 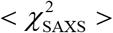 in the conformational space.

##### Summary of the final candidate models

All of the distributions of the rigid-body parameters in Fig. 5 have a single peak, suggesting that we were able to successfully narrow down the candidate models to a group with similar conformations. When the histone tails were not included in the models (step 3), two peaks appeared in the distribution of *dX* (Fig. S4), which corresponded to two groups of models with significantly different conformations at the atomic level (Fig. S5). When the histone tails were included, one group of the models gave smaller 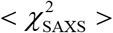 values and became more favorable, showing that inclusion of the histone tails contributed to the selection of the models. The model with the smallest 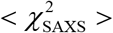 was included in this favorable group. The positions of the histone tails differed significantly between the two groups. As shown in Fig. 6a, the histone tails in the models in the group with smaller 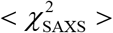 were observed more frequently at the interface between the two nucleosomes, or, more specifically, in the region where sequentially distant DNA sites came into close proximity. could be important for stabilizing the structure of OLDNs, as will be described later. On the other hand, tails in the other group were not observed in that region (Fig. 6b). Note that the model with the smallest 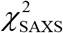 obtained when not considering the tails belonged to the unfavorable group (Fig. 6b).

**Figure 6.**
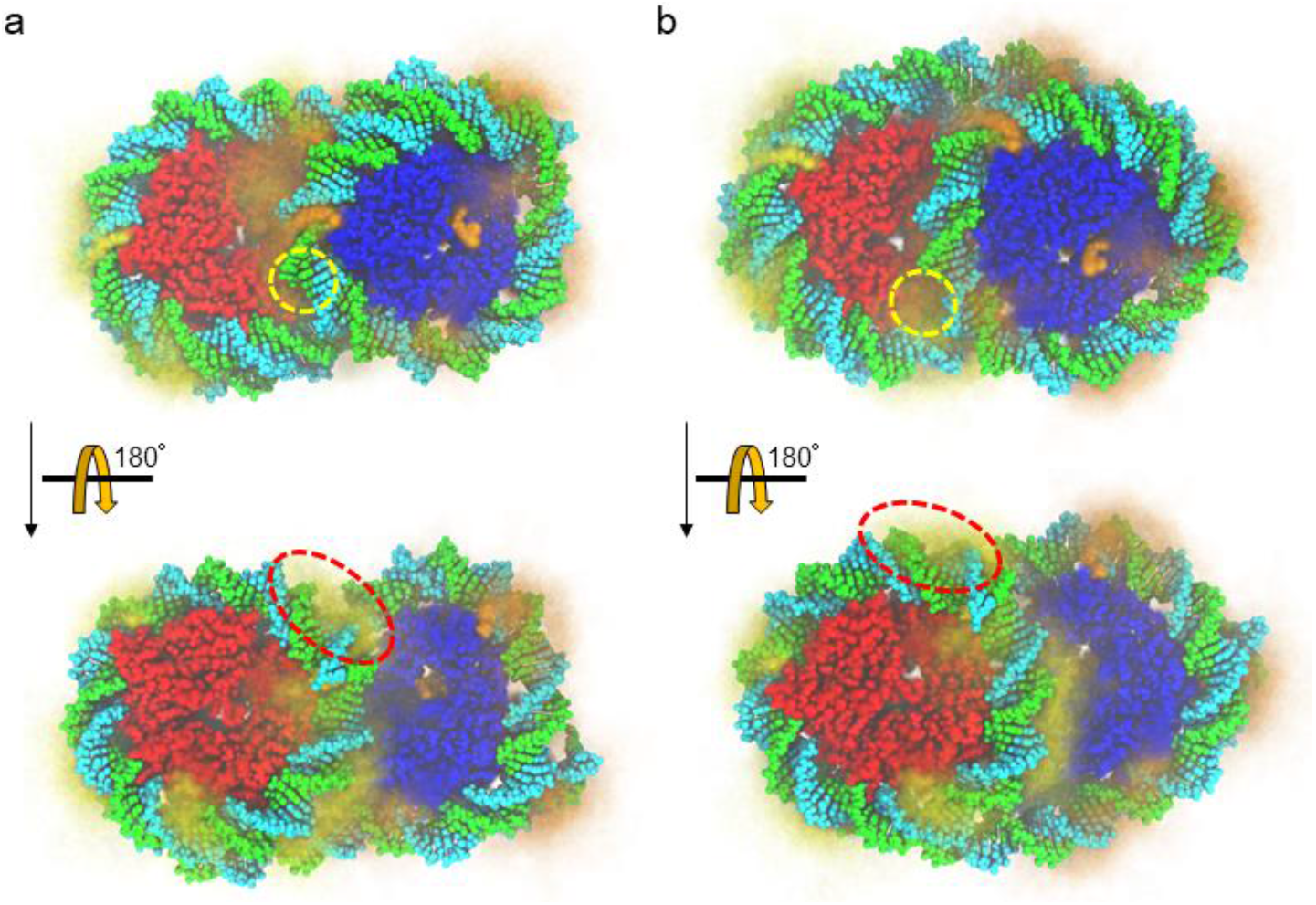
Atomic models of an OLDN. Multiple conformations of histone tails are illustrated by averaging 50 images of the model with histone tails in different conformations viewed from the same angle. The histone tails of the octamer and hexamer are colored in yellow and orange, respectively. Thick orange or yellow indicates that the histone tails are observed frequently in the area. (a) Model with the smallest 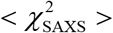. (b) Model with the smallest 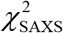 when the histone tails were not considered. Octameric core proteins are shown in red, hexameric in blue. The dashed circles in yellow and red respectively indicate the positions of one of the H4 and H3 histone tails in the octamer. Note that these circles are closer to both the hexasomal and octasomal DNA in (a) than in (b). The H4 histone tail indicated by the yellow circle in (a) is mostly beneath the hexasomal DNA.

Among the six rigid-body parameters, *dX*, *dρ*_*X*_, and *dρ*_*Y*_ distributions, suggesting that variations in these parameters had smaller effects on 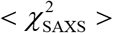, which is also apparent from the less steep slopes of the plots of 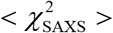 in Fig. 5. This result can be interpreted in two ways. One is that the OLDN is fluctuating in these directions; another is that it is difficult to differentiate the models deformed in these directions through SAXS analysis. To determine which interpretation is more likely, we carried out MD simulations, the results of which support the first interpretation, as will be described in step 6. The distributions of *dY*, *dZ*, and *dρ*_*Z*_ in Fig. 5 were narrow, suggesting that the variations in these parameters have larger effects on 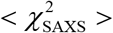.

Movie S1 shows the conformational changes in an OLDN when one of the six rigid body parameters in the model with the smallest 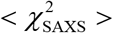 was forced to change while the other parameters kept unchanged as much as possible (see Supplementary text). When *dY* or *dρ*_*Z*_ was changed, however, other parameters also changed considerably (Fig. S10), which suggests these parameters are correlated with one another. This interpretation is described in detail in the Supplement.

##### Step 6: Model stability evaluated using MD simulations

To examine the stability of the models, we performed 10-ns-long all-atom MD simulations with explicit solvent models. From the final candidate models, we selected the 100 OLDN models with the smallest 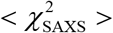 (< ∼6.75) for the simulations. The distributions of the rigid-body parameters of these models are shown as blue bars in Fig. 5. Among the 100 simulations, three failed during the initial energy minimizations due to bad positioning of the atoms in the models. For each of the remaining 97 models, the trajectory was saved and analyzed every 20 ps. The χ^2^ values and the six rigid-body parameters were computed for these conformations, as were the means and the standard deviations of the values in each trajectory.

Figure 7 shows the distributions of the 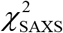 values for the initial conformations (or t=0) (a) and for all ∼50,000 conformations (b) during the simulations. The distribution of the latter conformations became wider toward both sides. It is noteworthy that the peak shifted toward a smaller 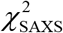 value, indicating that conformations that better fit the SAXS profile were sampled during the simulations. The standard deviations (SDs) of 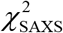 were generally small, and 25 of the 97 trajectories had SDs less than one (hereafter referred to as stable trajectories). One of the stable trajectories is shown in Movie S2. Figure S12 shows the distributions of the deviations in the six rigid-body parameters from the mean values in each trajectory. It is evident that *dX* and *dρ*_*X*_ had the largest variations among the translational (*dX*, *dY*, *dZ*) and rotational (*dρ*_*X*_, *dρ*_*Y*_, *dρ*_*Z*_) parameters, respectively. On the other hand, the variations in *dY* and *dρ*_*Z*_ were the smallest. These results are consistent with the distribution widths in the final models shown in Fig. 5, which indicates that the distribution widths obtained from the final models reflect the rigidity or flexibility of OLDNs in those directions.

**Figure 7.**
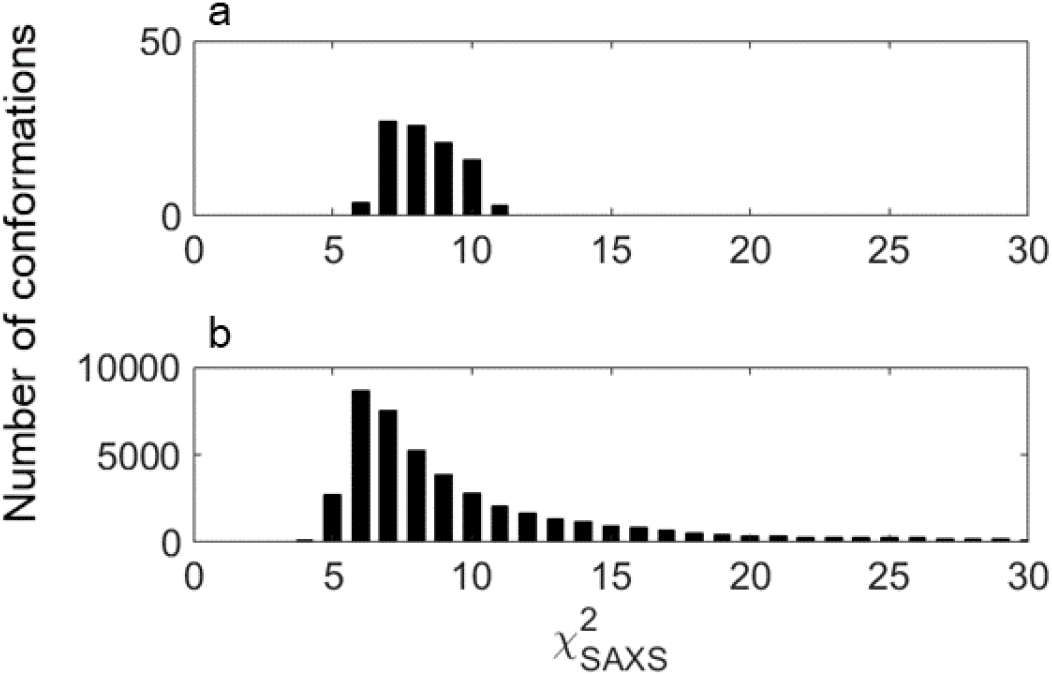
Distribution of 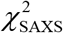 for the initial conformations (a) and for all (∼50,000) conformations (b) during the 97 trajectories of the MD simulations. The bin size was set to 1 in both panels.

Some trajectories exhibited structural instability, which yielded conformations with large 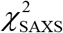 and widened the 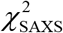 distribution (Fig. 7b). In such unstable trajectories, the two nucleosomes were often widely separated (Fig. 8b or Movie S3). Figure S13 shows the SD of 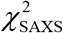 in each trajectory plotted against the mean value of the distance between the two nucleosomes, which was measured using the minimum interatomic distances between the hexasomal and octasomal DNA. The terminal 98 DNA base pairs wrapping the hexamer were regarded as the hexasomal DNA, while the other terminal 128 DNA base pairs were regarded as the octasomal DNA. Clearly, the deviation was large when the distance was more than 15 Å, and in all of the stable trajectories, the distances were less than 15 Å. We therefore concluded that the two nucleosomes comprising an OLDN were situated close to each other in the stable conformations, but the distance ranged from 2.5 to 15 Å.

**Figure 8.**
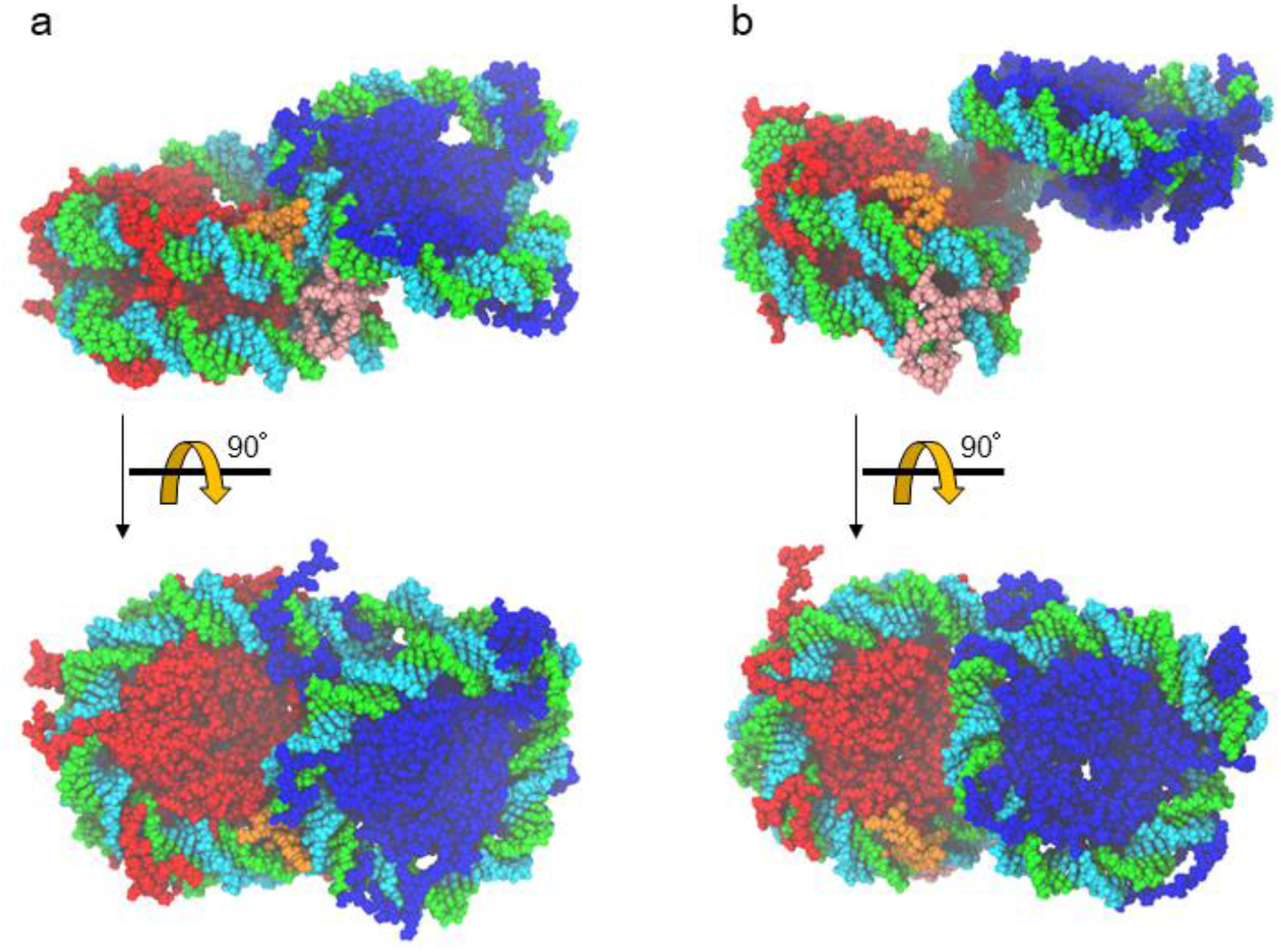
One of the conformations of OLDN in a stable trajectory where the standard deviation (SD) of 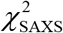 was less than 1 (a) and in a trajectory where the SD was large (b). Red: octameric proteins, blue: hexameric proteins, orange: one of the H4 histone tails in the octamer, pink: one of the H3 histone tails in the octamer.

We found that the conformations of the histone tails in the trajectories with small 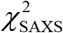 SDs clearly differed from those with larger SDs. In the small 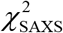 conformations, at least one of the histone tails of the octamer always bound simultaneously to both the octasomal and hexasomal DNA, acting as a bridge or glue between two DNA sites on opposite sides of the linker DNA (Fig. 8a or Movie S2). By contrast, no such histone tails were observed in the trajectories with large SDs (Fig. 8b or Movie S3). Because DNA is negatively charged, two DNA sites are unable to closely approach one another without the positively charged histone tails. In most trajectories, one of the H4 histone tails (orange in Fig. 8a) in the octamer served as this bridge. In some trajectories, one of the H3 histone tails (pink in Fig. 8a) in the octamer also served as a bridge.

## Conclusion

In this paper, we constructed atomic models of OLDNs in solution. Starting from the crystal structure, we first produced a library of atomic models with different conformations by deforming the DNA chain while keeping the structures of the histone proteins fixed. However, these models did not well reproduce the SAXS profiles. We therefore added the histone tails, which were invisible in the crystal structure. We then conducted repeated annealing MD simulations to generate a large number of different conformations of the tails. The addition of the histone tails improved the χ^2^ values for the SAXS profiles and enabled us to reduce the number of candidate models. We then used the SANS profiles for further refinement and selection of conformational candidates. The stability of the modelled structures was finally evaluated using MD simulations with explicit solvent models. These MD simulations showed that, in stable trajectories, the hexasomal and octasomal DNAs were close to one another, and that one or more histone tails were simultaneously bound to both DNA segments, which enabled the negatively charged DNA chains from the octamer and hexamer to be in close proximity.

## Author Contributions

A.Mat., M.S., H. Ku and H.Ko designed research, D.K. and A.O. prepared OLDN for SAXS and SANS, A.Mat., M.S., Z.L., A.Mar., L.P., R.I., H.Ku., and H.Ko. performed research, and A.Mat., M.S., Z.L., H.Ku., and H.Ko. wrote the manuscript.

## Acknowledgments

We thank Dr. Tomotaka Shimizu (KEK) for his help for SAXS experiments at BL10C KEK. We also thank Mr. Fumiya Adachi for his help to prepare OLDN for SAXS and SANS experiments. Molecular graphics were prepared using the software VMD (33). This work was supported by JSPS KAKENHI (18K06101 to A.Mat., JP18H05534, JP18H05229, JP16H01306 and JP15H02042 to M.S., JP17H01408 to H.Ku, JP25116003 to H.Ko, and JP18H05534 to H.Ku and H.Ko) and Platform Project for Supporting Drug Discovery and Life Science Research (BINDS) from AMED JP19am0101076 to H.Ku, and JP19am0101106 (support number 0363) to H.Ko. The SANS experiments at ILL were performed under proposals No. 8-03-824 and No.8-03-884 and the SAXS experiments at BL10C were also performed under proposal No. 2015G658.

## Supporting citations

Reference (34) appears in the Supporting Material.

